# Sulbactam-Durlobactam Plus Ceftriaxone Dosing and Novel Treatment Regimens for *Mycobacterium abscessus* Lung Disease

**DOI:** 10.1101/2025.08.05.668504

**Authors:** Sanjay Singh, Avneesh Shrivastava, Gunavanthi D Boorgula, Mary C Long, Brian Robbins, Pamela J McShane, Tawanda Gumbo, Shashikant Srivastava

## Abstract

**Background:** IDSA guideline-based therapy achieves sputum culture conversion rates in 20-34% of patients with *Mycobacterium abscessus* (MAB) lung disease (LD). Double-β-lactam combinations have been proposed to improve cure, based on time-kill curves.

**Methods:** We performed minimum inhibitory concentrations (MICs) experiments followed by hollow fiber system model of MAB-LD (HFS-MAB) exposure-effect studies with sulbactam-durlobactam administered every 8h (q8h), q12h, and q24h, to identify target exposures. Next, the sulbactam-durlobactam target exposure plus ceftriaxone was administered in the HFS-MAB inoculated with three different MAB isolates, as was the sulbactam-durlobactam-ceftriaxone combination with epetraborole and omadacycline (SDCEO). ***γ***-slopes (kill-speed) were calculated for all regimens. The minimal sulbactam-durlobactam clinical doses that achieved target exposure were identified using Monte Carlo experiments.

**Results:** Ceftriaxone reduced sulbactam-durlobactam MICs by 8-tube dilutions. In the HFS-MAB, sulbactam-durlobactam microbial kill and antimicrobial resistance were linked to % time concentration persists above MIC (%T_MIC_), with target exposure of 50%. Sulbactam-durlobactam killed 3.85 log_10_ CFU/mL below day 0 burden (*B_0_*) with regrowth. Sulbactam-durlobactam plus ceftriaxone killed without regrowth and demonstrated Bliss’ additivity. ***γ*** of bacterial population in >95% of virtual subjects were 2.28 (0.97-4.80) log_10_ CFU/mL/day for sulbactam-durlobactam-ceftriaxone and 2.91 (1.65-4.93) log_10_ CFU/mL/day for SDCEO. The optimal sulbactam-durlobactam dose co-administered with ceftriaxone was 2G q8h for creatinine clearance >90 mL/min, 2G q12h for 60-90 mL/min, 1G q12h for ≥30 to <60 mL/min, and 1G q24h for <30 mL/min.

**Conclusion:** Sulbactam-durlobactam-ceftriaxone achieved the highest microbial kill encountered so far in the HFS-MAB. Sulbactam-durlobactam-ceftriaxone should be tested as the backbone for novel treatment shortening regimens.

## INTRODUCTION

*Mycobacterium abscessus* (MAB) lung disease (LD) treated with guideline-based therapy (GBT) achieves sputum culture conversion in only 23-34% of patients [1]. The β-lactam antibiotics, cefoxitin and imipenem, are the backbone of GBT [2]. The poor success of GBT is because the MAB cell wall is relatively impermeable to antibiotics, and antimicrobial resistance (AMR) emergence is almost universal [3–5]. The MAB cell wall is composed of an outer mycolic acid layer, a middle arabinogalactan layer, and the inner peptidoglycan layer. Cell wall biogenesis is a 4-step process; the second step is peptidoglycan septum synthesis [6, 7]. β-lactams inhibit peptidoglycan synthesis by inactivating penicillin binding proteins (PBPs) such as D,D-transpeptidases, L,D-transpeptidases 1 to 5 [Ldt_Mab1_ to Ldt_Mab5_], the bifunctional transglycosylase, and D,D-transpeptidase PonA1. D,D-transpeptidases catalyze the formation of 4,3-cross-links of peptidoglycan stem peptides, while Ldt_Mab1-5_ catalyze the formation of 3,3-cross-links [8]. PonA1 transglycosylase links repeating disaccharide units of the glycan backbone while the transpeptidase crosslinks pentapeptide tails [6, 7]. β-lactams also inhibit D,D-carboxypeptidases (D,D-c), which converts peptidoglycan pentapeptides to tetrapeptides to be used by Ldts [9]. A potential therapeutic strategy, termed “target redundancy”, would involve simultaneous targeting of all the PBPs [10].

*MAB* harbor the β-lactamase, Bla_Mab_, and therefore, β-lactamase inhibitors (BLI) such as sulbactam (a β-lactam) and durlobactam (a diazabicyclooctane) represent an attractive strategy to combat AMR [11, 12]. The BLIs and β-lactams have varying affinities for Bla_Mab_ and PBPs [8, 11–15]. In a previous study, ceftriaxone and cefotaxime inactivated PonA2, PonA1, and D,D-c PbpA at low concentrations, while ceftazidime and cefoxitin inactivated the same at intermediate concentrations, but the BLIs, such as sulbactam, required higher concentrations [15]. However, durlobactam inactivated Bla_Mab_ and all Ldt_Mab_ (except Ldt_Mab3_) more potently than avibactam [16]. Therefore, target redundancy strategy has been proposed for durlobactam, sulbactam, imipenem, cefuroxime, ceftaroline, and ceftazidime [8, 10, 17–20].

Here, we tested ceftriaxone plus sulbactam-durlobactam against four isolates of MAB. We also tested experimental combinations with epetraborole, and omadacycline, with reported activity against MAB [21–23]. Since the shape of the concentration-time curve is crucial to β-lactam efficacy, and an oscillating stressor (concentration-time profiles with repetitive dosing) more easily generates AMR, we conducted a series of studies in which we mimicked intrapulmonary pharmacokinetics (PKs) of drugs in the hollow fiber system model of MAB-LD (HFS-MAB) [4, 24–36]. The HFS-MAB offers repetitive sampling, thus enabling tracking of MAB burdens (CFU/mL) treated with different drug exposures over time. The HFS-MAB fits the “FDA Roadmap to Reducing Animal Testing in Preclinical Safety Studies” approach https://www.fda.gov/media/186092/download?attachment and the NIH initiative to phase out animal models for therapeutics for human diseases, as do the tandem *in silico* translations [37, 38].

## METHODS

### Bacterial isolates and materials

The reference laboratory isolate was purchased from the American Type Culture Collection (ATCC; ATCC#19977). Sixty-three clinical isolates for MIC testing (including two for HFS-MAB experiments), 39 were *MAB subspecies abscessus*, 21 *MAB subsp. massilliense*, and 3 *MAB subsp. bolletii*.

### Minimal Inhibitory concentrations and time-kill studies

Minimal inhibitory concentrations (MICs) were tested as described in detail **in Supplementary Methods.**

### HFS-MAB exposure-effect

The construction and design of the HFS-MAB have been extensively described in the literature [4, 24–30]. Here, we combined an exposure-effect study with a dose-scheduling design, using ATCC#19977. Drugs were infused over 14 days to mimic the intrapulmonary PKs following human equivalent doses at every X hours (QXh) as follows: 0, 3G Q24h, 1.5G Q12h, 1G Q8h, 4.5G Q24h, 3G Q12h, 1.5G Q8h, 6G Q24h, 3G Q12h, 2G Q8h, and 3G Q8h. The drug exposures and sampling details are shown in the **Supplementary Methods and Supplementary Table S1.**

### HFS-MAB double-β-lactam-β-lactamase inhibitor combination in three more MAB isolates

For this set of experiments, we chose ceftriaxone based on the rationale in **Supplementary Methods** and **Supplementary Table S1** [15, 39–41]. We examined the effect of sulbactam-durlobactam at target exposures alone, ceftriaxone alone, and sulbactam-durlobactam- ceftriaxone versus non-treated HFS-MAB replicates, using two MAB clinical isolates in the HFS- MAB. In a third HFS-MAB isolate, we compared the sulbactam-durlobactam-ceftriaxone regimen to combinations of sulbactam-durlobactam-ceftriaxone plus omadacycline and epetraborole (SDCEO), and another regimen in which omadacycline was replaced by minocycline (SDCEM). Epetraborole and omadacycline were chosen because they have the best kill rates below *B_0_* in the HFS-MAB, based on our meta-analysis (*Srivastava and Gumbo. Submitted*). Sulbactam-durlobactam was administered Q12h, while ceftriaxone was delivered Q24h. The target concentrations, sampling strategy for PKs and bacterial burden for all three isolates is described in **Supplementary Methods** and **Supplementary Table S1.**

### Mathematical modeling

First, we modeled CFU/mL using the inhibitory sigmoid E_max_ model for log_10_ CFU/mL versus exposure for microbial kill, while for resistance we used the quadratic function of Gumbo et al, for each sampling day [3, 42]. We used corrected Akaike Information criteria to identify the sulbactam-durlobactam PK/PD driver (relevant to dosing schedule) for microbial kill and AMR. Second, data for drug combinations were analyzed based on a system of ordinary differential equations (ODEs), we developed for other mycobacteria in the hollow fiber system and patients, which we previously adapted for MAB [40, 43]. This allowed us to calculate the bacteria time-to-extinction (TTE), which is the time to achieve MAB burden of 10^-2^ CFU/mL, as described in the past [40, 43].

### Monte Carlo experiments (MCE)

MCE is a tool for *in silico* dose-ranging and dose scheduling experiments, to identify the drug dose for use in the clinic. MCEs were performed as described in detail in the **Supplementary Methods**. The population PK parameters used were those of Cammarata et al, which account for between-patient PK variability [44]. The probability of target attainment (PTA) is the proportion of patients who will achieve target exposure at each MIC, given the PK variability. The cumulative fraction of response (CFR) to each dose, which is the proportion of all patients in which dose achieves target exposures across the entire MIC distribution, was calculated for each of the four creatinine clearance categories. These were creatinine clearance (i) >90 mL/min which is normal renal function, (ii) 60-90 mL/min which is mild renal dysfunction, (iii) ≥30 to <60 mL/min moderate renal dysfunction, and (iv) <30 mL/min which is severe. We also examined the dose of the co-administered ceftriaxone; the PK model used was that we have published elsewhere for MAC-LD [40]. Since durlobactam and sulbactam will have different PK/PD target exposures, we chose the optimal dose as that at which the CFR was ≥90% for both sulbactam and durlobactam [40].

### Quality score of the HFS-MAB studies

The reproducibility and rigor applied to HFS-MAB were measured using our novel quality score [45]. HFS-MAB must be of high quality, especially since results are used for identifying clinical dosing. We applied this tool to studies performed here.

## RESULTS

### MICs

The MICs of sulbactam-durlobactam alone, ceftriaxone alone, and sulbactam-durlobactam plus ceftriaxone for each MAB clinical isolate are listed in **Supplementary Table S2** and summarized in **Figure 1A**. The sulbactam-durlobactam MIC_50_ was 32 mg/L, and the MIC_90_ was 64 mg/L. When co-incubated with a fixed ceftriaxone concentration of 256 mg/L, the sulbactam-durlobactam MICs were mostly <0.125 mg/L (for calculation purposes categorized as 0.06 mg/L), leading to MIC_50_ of 0.06 mg/L and an MIC_90_ of 8 mg/L. Thus, ceftriaxone reduced sulbactam-durlobactam MICs by a median of 8-tube dilutions (p<0.001).

**Figure 1.**
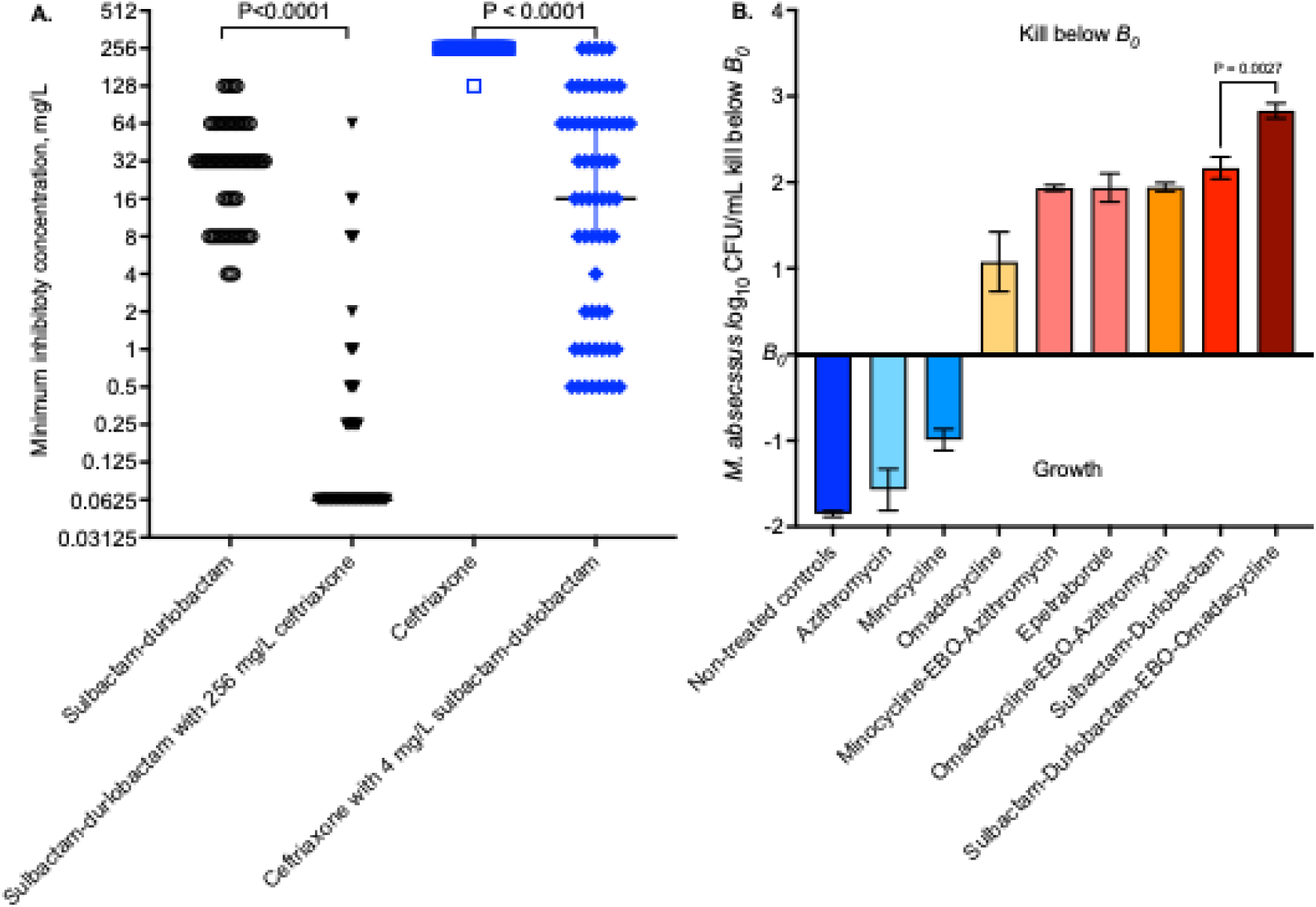
Minimal inhibitory concentrations and time-kill with different drugs and combinations. All assays were performed in triplicate. **A.** MICs were performed for 63 MAB isolates, under 4 conditions: sulbactam-durlobactam, sulbactam-durlobactam with 256 mg/L ceftriaxone (trio), ceftriaxone, and ceftriaxone plus 4 mg/L sulbactam-durlobactam (trio). Shown are the reductions in MIC for trios. **B.** Microbial kill below day 0 with drug alone and in combination after 72h of co-incubation.

The ceftriaxone MICs are shown in **Figure 1A**. The ceftriaxone MIC_50_ and MIC_90_ were >128 mg/L (graphed as 256 mg/L). However, in the presence of a fixed sulbactam-durlobactam concentration of 4 mg/L, the ceftriaxone MIC_50_ was 1 mg/L and MIC_90_ was 64 mg/L. The ceftriaxone MICs in the presence of sulbactam-durlobactam correlated with sulbactam-durlobactam MICs (p<0.001) with a Pearson r of 0.42 (95% confidence interval [CI] 0.18-0.61). This means that the ceftriaxone MICs were partially driven by the sulbactam-durlobactam effective concentration. In addition, even though all sulbactam-durlobactam MIC_50_ of 32 mg/L was higher than 4 mg/L used in the assay, sulbactam-durlobactam still reduced ceftriaxone MICs. In short, sulbactam-durlobactam reduced ceftriaxone by 4-tube dilutions (p<0.001).

### Time-kill studies

**Figure 1B** shows the effects of different drugs, including sulbactam-durlobactam, with 72 h of co-incubation with MAB. Azithromycin and minocycline did not kill below the day 0 burden (B_0_). Sulbactam-durlobactam duo killed 2.17±0.13 log_10_ CFU/mL below *B_0_,* better than the macrolide and tetracycline-containing combinations. Sulbactam-durlobactam combination with epetraborole and omadacycline killed 2.83±0.09 log_10_ CFU/mL below *B_0_*, better than sulbactam-durlobactam duo.

### Sulbactam-durlobactam exposure-effect and dose scheduling study in the HFS-MAB

The observed drug concentration-time profiles and PKs are shown in supplementary data section, and for sulbactam in **Supplementary Figure S1A-D** while for durlobactam are shown in **Supplementary Figure S2A-D.** The PK modeled data for sulbactam and durlobactam were used to calculate AUC/MIC ratios, peak concentration to MIC ratio, and time concentration persists above MIC (%T_MIC_); the MAB ATCC#19977 inoculated in HFS-MAB had an MIC of 4 mg/L for sulbactam-durlobactam (1:1 combination).

The day-to-day sampling revealed changes in bacterial burden shown in **Figure 2A**. The bacterial burden on day 0 (*B_0_*) was 4.88 log_10_ CFU/mL. In non-treated HFS-MAB, bacterial burden grew to 8.46 log_10_ CFU/mL on day 14. The highest microbial kill below *B_0_* occurred on day 3. After day 4 the bacterial burden began to increase. Since sulbactam degrades at 37°C when supplemented in agar, we could not quantify the drug-resistant CFU/mL. However, MICs of samples from each HFS-MAB unit on day 14 demonstrate that while the MIC did not change in non-treated controls (4 mg/L), it increased by one-tube dilution (8 mg/L) in the lowest exposure and three-tube dilutions (32 mg/L) in the rest of the sulbactam-durlobactam-treated HFS-MAB units.

**Figure 2.**
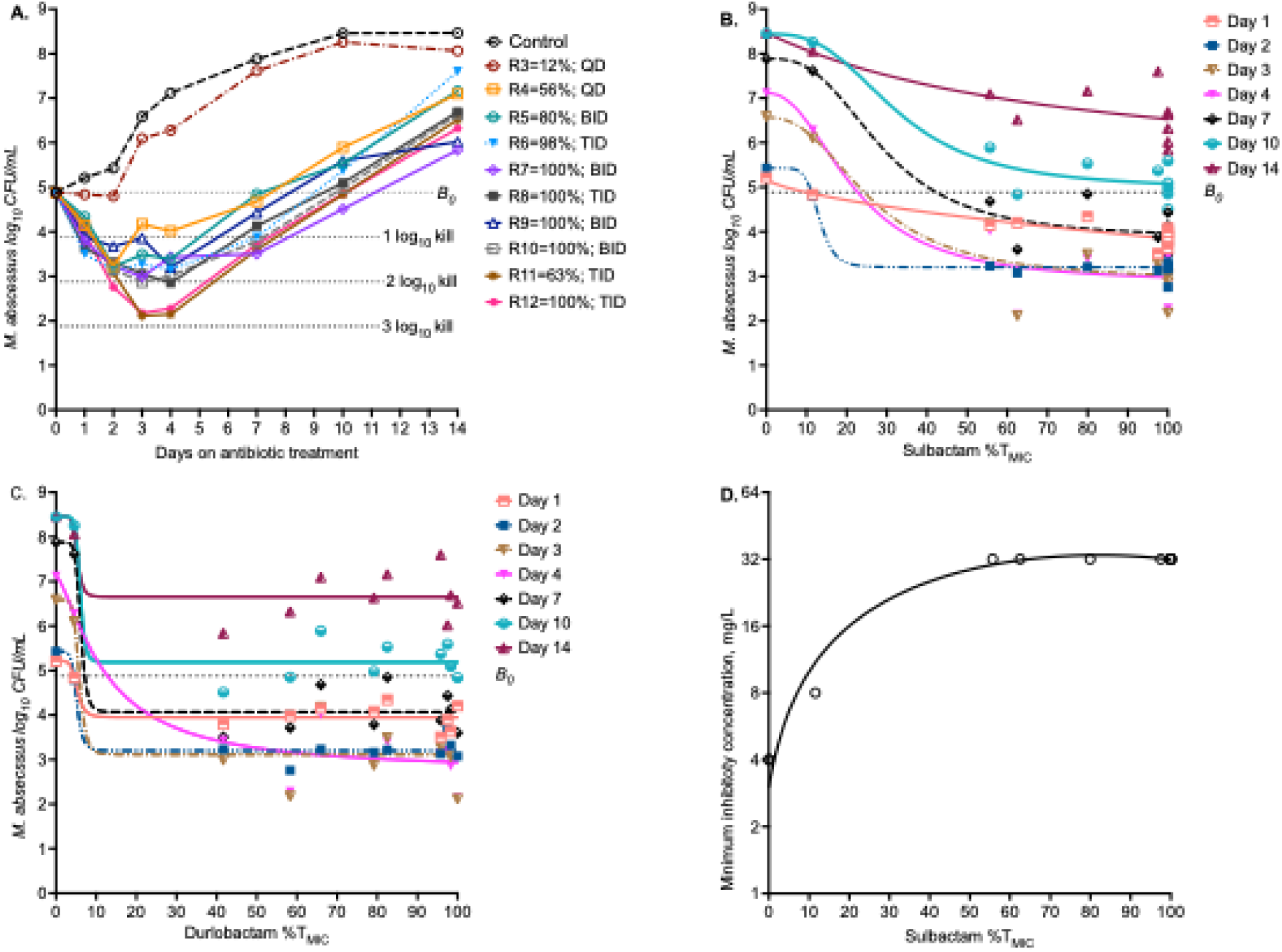
Sulbactam-durlobactam pharmacokinetics/pharmacodynamics in the HFS-MAB. **A.** Changes in bacterial burden between sampling days on different treatment regimens, shown with percentage of dosing interval sulbactam concentrations stayed above MIC. All treatment regimens demonstrated a biphasic effect. **B.** Inhibitory sigmoid E_max_ curves based on sulbactam exposure, for the different sampling days. **C.** Inhibitory sigmoid E_max_ curves based on durlobactam exposure, for the different sampling day, shows that steep portions of the curves were between 0 and 10% exposures. **D.** Quadratic function for MICs versus sulbactam exposure. (Sul/Dur, sulbactam/durlobactam)

The PK/PD driver for microbial kill for sulbactam was %T_MIC_, as shown in **Supplementary Table S3**. For durlobactam microbial kill %T_MIC_ was also the driver, except on day 2 when AUC_0-24_/MIC was the PK/PD driver. The relationship between sulbactam %T_MIC_ and total bacterial burden is shown in **Figure 2B**, while that for durlobactam is shown in **Figure 2C**. The inhibitory sigmoid E_max_ parameter estimates are shown in **Table 1**, as are the target exposures (EC_80_) for each day. The mean sulbactam target exposure was %T_MIC_ of 49.95 ± 8.43% or ∼50%, while that for durlobactam was 10.5±3.8%. When examined using microbial kill below *B_0_*, the maximal kill from the model was 3.85±1.3 log_10_ CFU/mL, on day 4. As regards to AMR, the relationship between sulbactam %T_MIC_ and MIC on day 14 is shown in **Figure 2D** (r^2^>0.99), associated with an Akaike information criteria score of 17.16 as compared to 51.35 for AUC_0-24_/MIC and 51.04 for C_max_/MIC. This means %T_MIC_ is also PK/PD driver for sulbactam-durlobactam AMR. Results were similar for durlobactam versus MIC (r^2^=0.98).

**Table 1.**
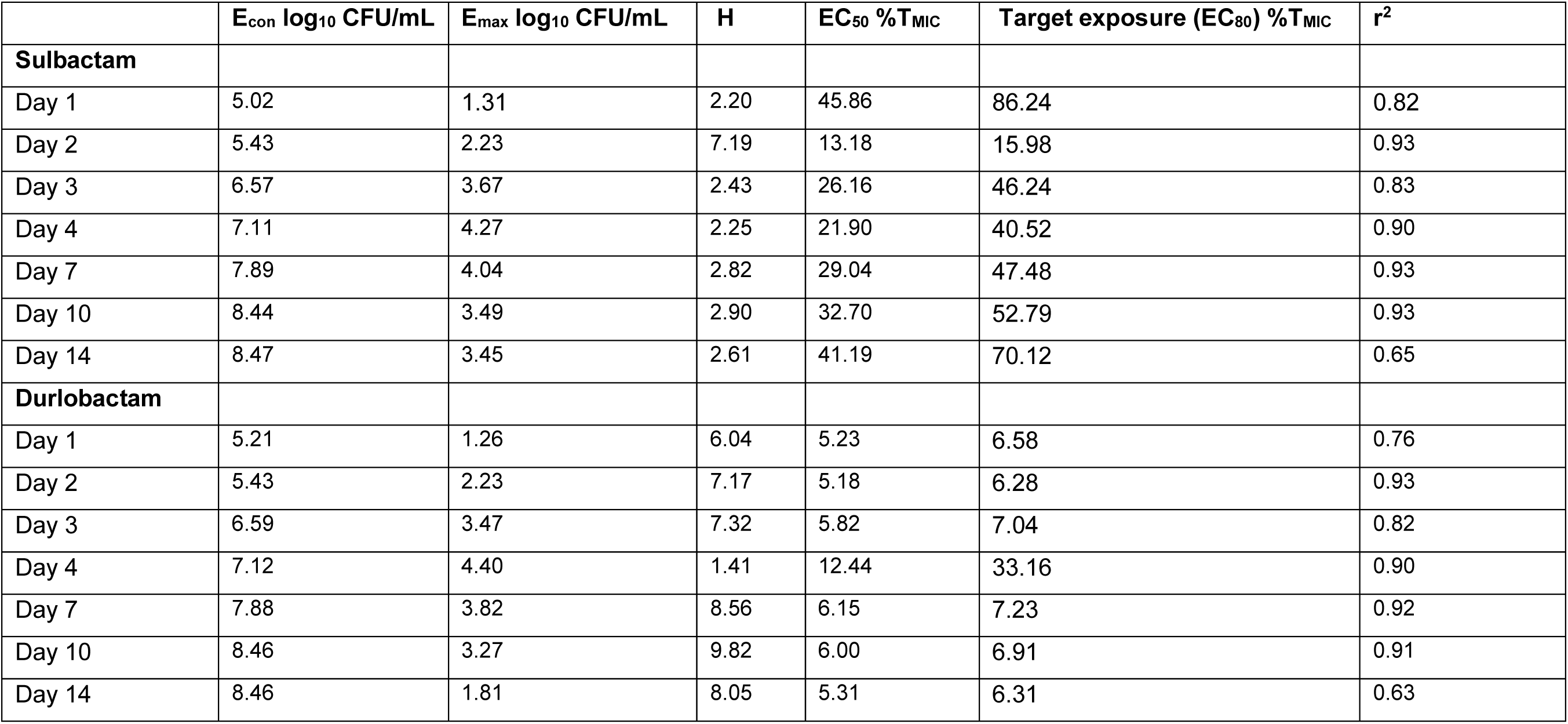
Inhibitory Sigmoid E_max_ Model Parameter Estimates for sulbactam and durlobactam on different sampling days.

### Combination therapy studies and generalization using 3 clinical isolates in the HFS-MAB

The three MAB isolates had MICs shown in **Supplementary Table S4**. The drug concentration-time profiles measured in the combination therapy studies were as shown in **Supplementary Figure S3.** The exposures achieved for each drug are shown in **Supplementary Table S5.**

Given that each MAB isolate achieved different *B_0_*, and non-treated systems achieve different bacterial burdens (*B_x,t_*) from each other on each sampling day (*t*) due to differences in growth rates, we modeled effect as kill below *B_0_*, with results shown in **Figure 3A**. Kill below *B_0_* is a positive number and growth above *B_0_*is a negative number. **Figure 3A** shows that while ceftriaxone monotherapy had some effect, the effect was minimal. Sulbactam-durlobactam duo killed MAB better than ceftriaxone but was followed by rebound growth. In addition, **Figure 3A** also shows that the sulbactam-durlobactam-ceftriaxone combination trio was more effective and killed about 2 log_10_ below *B_0_* and then held the bacterial burden constant without rebound growth. Furthermore, **Figure 3A** shows that all the sulbactam-durlobactam-ceftriaxone combinations with epetraborole or tetracyclines [omadacycline (SDCEO) versus minocycline (SDCEM)] killed more than sulbactam-durlobactam-ceftriaxone trio. In **Figure 3B**, we compared the observed sulbactam-durlobactam-ceftriaxone trio kill below *B_0_* to the sum of ceftriaxone monotherapy plus sulbactam-durlobactam duo (theoretical Bliss-type additivity). The observed trio effect fell within the 95% confidence interval of the theoretical additivity. This means that the effects of sulbactam-durlobactam and ceftriaxone were additive, and not antagonistic. As regards to resistance, the MICs on day 14, at the end of the experiments, are shown in **Supplementary Results and Table S4**.

**Figure 3.**
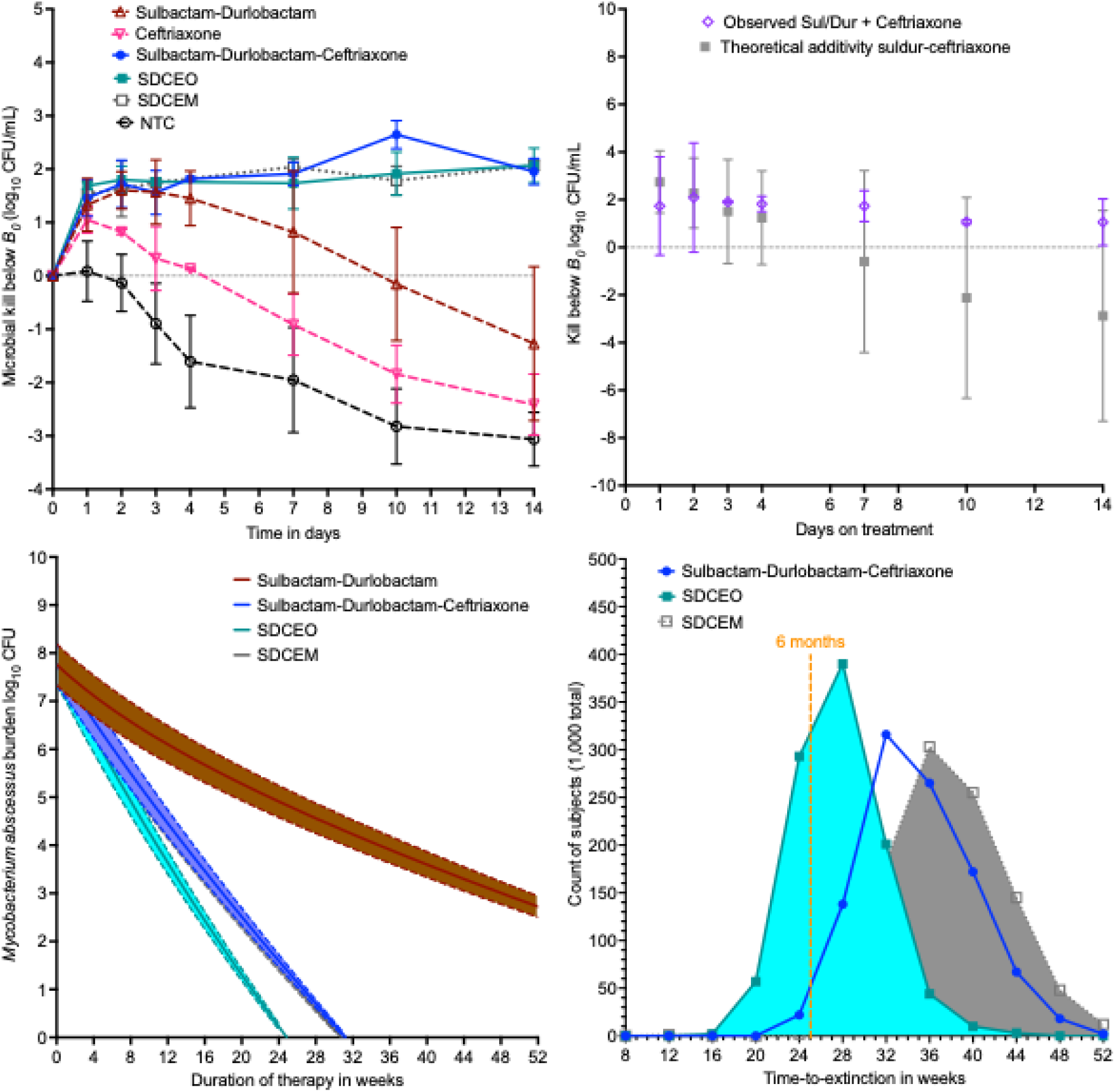
Combination therapy pharmacodynamics and bacterial population extinction. SDCEM is Sulbactam-durlobactam-ceftriaxone plus epetraborole plus minocycline; SDCEO is sulbactam-durlobactam-ceftriaxone plus epetraborole plus omadacycline; NTC are non-treated controls. **A.** Kill below day 0 bacterial burden (*B_0_*) for the different regimens. Symbols show mean value, and error bars standard deviation. **B.** Error bars are 95% confidence intervals; symbols are mean values. **C.** Modeling of bacterial burden decline till extinction, showing the median and 95% credible intervals. Time-to-extinction is shown for those regimens that achieved that at or before 52 weeks.

### ***γ***, time-to-extinction, and time to cure

The results of all isolates (including from ATCC#19977) were combined and co-modeled for ***γ*** for each of the six regimens, with results shown in **Figure 3C**, and **Table 2**. **Figure 3C** shows that the model sampled from a *B_0_* between 7.0 and 8.0 CFU. ***γ*** is the speed of kill and ranked by this parameter the regimen with the fastest kill was SDCEO, followed by SDCEM. Time-to-extinction is shown in **Figure 3D**. The % of virtual subjects with MAB population extinction at 52 weeks (1 year) was 0.9% ceftriaxone monotherapy, 38.4% for sulbactam-durlobactam, 99.9% for sulbactam-durlobactam-ceftriaxone, 98.9% for SDCEM, and >99.9% for SDCEO. For sulbactam-durlobactam-ceftriaxone the mean time-to-extinction was 34.43 (95%CI: 34.1-34.75) weeks, SDCEM 37.8 (95%CI: 37.39-38.21) weeks, and for SDCEO 27.15 (95%CI: 26.91-27.4) weeks.

**Table 2.** Ordinary Differential Equation Parameter Estimates and 95% Credible Intervals.

| | $B_0 \log_{10}$<br>CFU/cartridge | $r \log_{10}$<br>CFU/ml/day | $K_{\max} \log_{10}$<br>CFU/catridge | $\gamma$<br>$\log_{10}$ CFU/ml/day | Time-to-extinction in 95%<br>of subjects in weeks |
| --- | --- | --- | --- | --- | --- |
| Sulbactam-durlobactam | 6.14 (5.22-6.32) | 0.18 (0.05-0.32) | 10.15 (8.94-26.5) | 1.36 (0.74-3.06) | >52 |
| Ceftriaxone | 5.56 (5.13-5.96) | 0.17 (0.11-0.25) | 9.59 (8.93-10.72) | 0.94 (0-2.81) | >52 |
| Sulbactam-durlobactam + ceftriaxone | 6.62 (6.01-7.21) | 0.16 (0.07-0.29) | 10.37 (9.48-13.11) | 2.28 (0.97-4.80) | 43.75 (31.12-57.92) |
| Sulbactam-durlobactam + ceftriaxone +<br>epetraborole + omadacycline (SDCEO) | 8.03 (7.74-8.31) | 0.12 (0.07-0.18) | 11.35 (10.74-12.69) | 2.91 (1.65-4.93) | 33.75 (23.55-47.5) |
| Sulbactam-durlobactam + ceftriaxone-<br>epetraborole + minocycline (SDCEM) | 8.03 (7.74-8.31) | 0.12 (0.07-0.18) | 11.35 (10.74-12.69) | 2.30 (0.80-4.83) | 46.00 (33.68-61.36) |
| **ALL ISOLATES** | **6.87 (6.3-7.41)** | **0.15 (0.06-0.27)** | **10.57 (9.69-13.33)** | - | - |

$B_0$ is pretreatment bacterial burden; $K_{\max}$ is carrying capacity which is the maximum possible bacterial population carried by hollow fiber system cartridge; $\gamma$ is kill rate or slope from ordinary differential equation.

As regards to sulbactam-durlobactam or ceftriaxone, the mean time-to-extinction was >52 weeks. When analysis was performed for time-to-extinction for >95% of the virtual subjects (i.e., time-to-cure), results were as shown in **Table 2**. SDCEO was predicted to achieve that in 33.75 weeks, while sulbactam-durlobactam-ceftriaxone was predicted to achieve cure at 43.75 weeks.

### Probability of target attainment (PTA) for clinical dose selection

MCE were performed in a total of 480,000 virtual subjects for sulbactam and 480,000 for durlobactam, across a range of creatinine clearances. The PK output in the virtual subjects was compared to that used in the domain of input of the MCE in **Supplementary Table S6**. The MCE thus recapitulated observed PKs well.

The sulbactam probability of target attainment (PTA) is the proportion of patients achieving the target exposure of %T_MIC_ of 50% in the lung at each MIC, for the different creatinine clearance categories were as shown in **Figure 4**. First, the PTA improved as renal dysfunction worsened, as shown from considering the PTA for the lowest dose of 1G Q24h at the MIC of 2 mg/L, which was 1% with normal renal function (**Figure 4A**), 9% with mild (**Figure 4B**), 40% with moderate **(Figure 4C)**, and 96% with severe renal dysfunction (**Figure 4D**). Second, the two MIC distributions for sulbactam-durlobactam without and with fixed ceftriaxone concentration of 256 mg/L are also shown in **Figure 4**. The Q8h dosing schedule achieved sulbactam PTA ≥90% above the MIC_90_ of MIC distribution sulbactam-durlobactam with ceftriaxone versus at the MIC_10_ of MIC without ceftriaxone, for normal renal function. This means that renal function, dosing schedule, and use of ceftriaxone were important drivers for achievement of >90% PTA for sulbactam. The PTAs for durlobactam are shown in supplementary results, and **Supplementary Figure S4A-D.**

**Figure 4.**
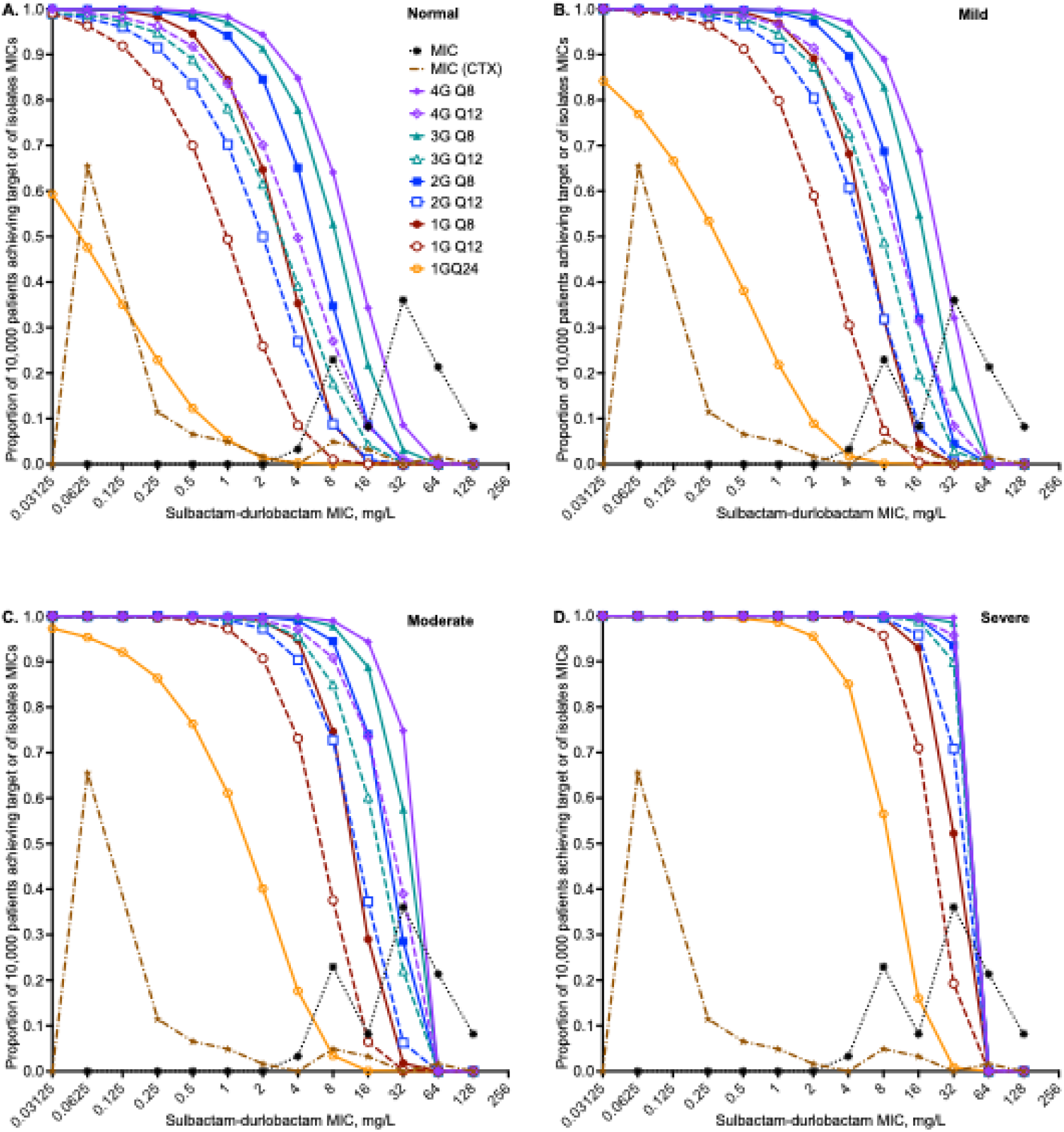
Probability of target attainment for sulbactam. MIC (CTX) refers to sulbactam-durlobactam MIC with 256 mg/L of ceftriaxone. Every 8h dosing (q8h) is shown with solid symbols and lines, every 12h (q12h) open symbols and hatched lines, and the 1G q24h dose is the hexagon. **A.** Probability of target attainment (PTA) in 10,000 virtual subjects with creatinine clearance of >90 mL/min. **B.** PTA in 10,000 virtual subjects with creatinine clearance of 60-90 mL/min, which is mild renal dysfunction. **C.** PTA in 10,000 virtual subjects with creatinine clearance of ≥30 to <60 mL/min**. D**. PTA in 10,000 virtual subjects with creatinine clearance <30 mL/min.

The cumulative fraction of response (CFR), which is the proportion of patients that achieve target exposures when all MICs are summated, is shown by renal function in **Figure 5A-D**. The lowest dose that achieves CFR in >90% of virtual patients is the best dose for the clinic (i.e., optimal). Since durlobactam CFR was achieved at much lower doses compared to sulbactam, the sulbactam CFR values were used to define the optimal sulbactam-durlobactam dose. First, **Figure 5** shows that under all the different degrees of renal function, the co-administration of sulbactam-durlobactam with ceftriaxone achieved the target in a larger proportion of patients compared to when there was no co-administration of ceftriaxone. Indeed, all the doses based on sulbactam-durlobactam without ceftriaxone failed to achieve the target in 90% of patients even up to 4G Q8h (12g per day. Second, Figure 5 shows that the lowest durlobactam dose of 1G Q24h achieved a CFR >90% for all regardless of renal function. Thus, the lowest doses in which both sulbactam and durlobactam achieved CFR ≥90% were chosen based on sulbactam CFRs.

**Figure 5.**
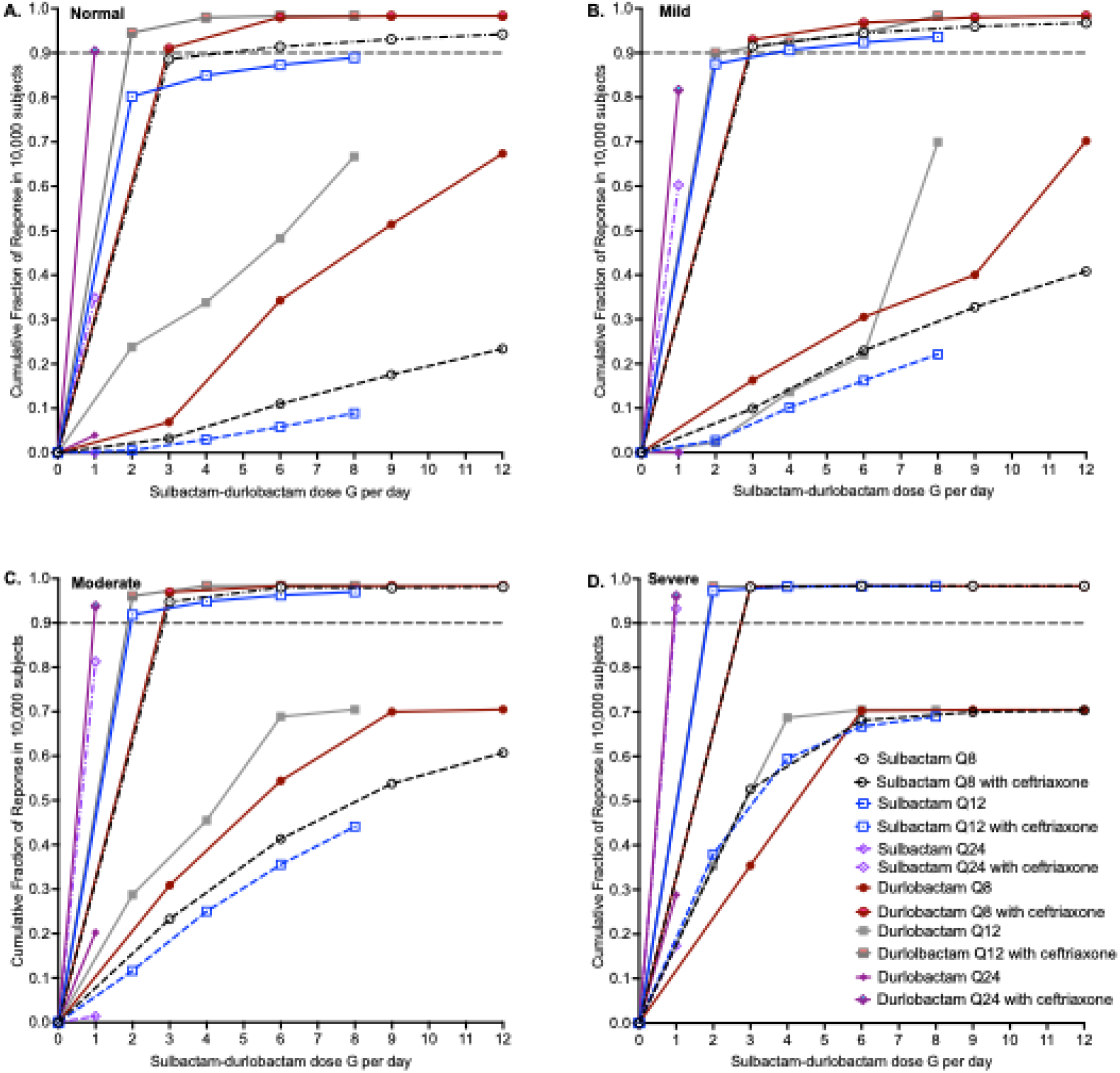
Cumulative Fraction of Response for both sulbactam and durlobactam. Cumulative fraction of response (CFR) in patients for both sulbactam target exposure (%T_MIC_=50%) and durlobactam exposure target (%T_MIC_=10.5%). Figure legends are shown in panel D. Open symbols and hatched lines are for sulbactam, while closed symbols and solid lines are for durlobactam. Circles are for Q8h dosing, squares Q12h, and diamonds Q24h. Pairs with and without ceftriaxone are color coordinated. The proportion of 90% of virtual patients achieving target attainment across all MICs is shown by the grey hatched line. **A.** CFR in 90,000 virtual patients with creatinine clearance >90 mL/min. **B.** CFR in 90,000 virtual patients with creatinine clearance 60-90 mL/min. **C.** CFR in 90,000 virtual patients with creatinine clearance ≥30 to <60 mL/min. **D.** CFR in 90,000 virtual patients with creatinine clearance <30 mL/min.

In patients with normal renal function, 2G Q8h was the optimal dose (**Figure 5A**), no sulbactam Q12h or Q24h doses achieved a CFR >90%. In **Figure 5B**, at a creatinine clearance of 60-90 mL/min, the sulbactam CFR were >90% at the lowest doses of 1G Q8h or 2G Q12h. At a creatinine clearance of ≥30 to <60 mL/min (**Figure 5C**), sulbactam 1G Q12h was the optimal dose. With creatinine clearance less than 30 ml/min (**Figure 5D**), 1G Q24h was optimal the optimal. This dosing is summarized in **Supplementary Table S7** for clinical use.

### Quality score of the study

Our study had a quality score of 21 out of a total possible of 24. This means the reported studies are categorized as high quality.

## DISCUSSION

First, in patients’ lung cavities, pretreatment burden (*B_0_*) ranges between 7.0-8.0 log_10_ CFU/lung, and is the major determinant of therapy outcomes, consistent with the inoculum effect [46, 47]. Therefore, drugs for the target redundancy strategy must be chosen based on how extensively they kill MAB below *B_0_* [26, 28]. In addition, ceftriaxone has an 8h half-life, and sulbactam-durlobactam have half-life of 2.5h, which could lead to once or twice a day dosing, more practical than imipenem, ceftaroline, and cefoxitin, dosed four to six times a day [26, 28, 30].

Second, ceftriaxone and sulbactam-durlobactam reduced each other’s MICs by multiple tube dilutions. This means that it will be necessary to develop MIC assays for the “double β-lactam” combinations intended for use, for calculation of MIC_50_ and MIC_90_, as well as for epidemiologic cut-off values, in place of the classic approach of examining each drug alone. This can be accomplished in MIC assays by adding a fixed concentration of one drug (either sulbactam-durlobactam or ceftriaxone) throughout the wells and then varying the concentration of companion drug through a range of concentration series.

Third, time-kill curves studies have shown that sulbactam-durlobactam alone and in combination with ceftaroline, cefuroxime, cefoxitin and imipenem, killed more than GBT [16, 48]. This was confirmed by our own time kill curves in **Figure 1B**. However, time-kill studies have no straightforward translation to clinical dosing schedules, and cannot be used to identify %T_MIC_ target exposures, because %T_MIC_ is either 0 or 100% in time-kill assays. The HFS-MAB mimicks intrapulmonary PKs in order to identify (1) the target exposure and hence optimal clinical dose, and (2) optimal dose-schedule that drives microbial kill and AMR. Furthermore, ceftriaxone was demonstrated to be additive to sulbactam-durlobactam as regards to microbial kill on all sampling days in the HFS-MAB, preventing rebound growth.

Fourth, what do the HFS-MAB findings mean regarding sulbactam-durlobactam dosing strategy in the clinic? In the treatment of Gram-negative bacillary sepsis, sulbactam-durlobactam is administered with a 3h infusion at a schedule of every 4h, which would be impractical for administration over many months in patients with MAB-LD. In the MCEs the inclusion of ceftriaxone shifted MICs to a range that could be covered at dosing schedules of three-times a day or less. The sulbactam-durlobactam doses and dose schedules, based on renal function, are shown in **Supplementary Table S7**, for use by clinicians. The mathematical modeling gave us insights regarding duration of therapy. Use of ***γ***-modeling allows for pooling data from different experiments, and ranking drugs by how fast they kill MAB. Moreover, since sulbactam- durlobactam demonstrated biologic activity against all four MAB isolates in the HFS-MAB, efficacy can be generalized to account for heterogeneity of MAB response in different patients.

Finally, we made a methodological contribution to MCEs for the two drugs in a fixed-dose combination, including a third drug. Sulbactam had a different target exposure from durlobactam, so the question was which drug’s PTA and CFR to use. Renal function had differential effects of sulbactam compared to durlobactam. Should the choice be made based on the drug with better kill below *B_0_* (durlobactam)? Here we introduced methods how to use both drugs in the combination. We also explored effect of a third drug, ceftriaxone, which has three effects, (1) on MIC distribution, (2) on extent of microbial kill and rebound, and (3) on target exposure and dose-schedule. These concepts inform us of how we should explore combination therapy in MCEs in general theraputics.

There are some limitations to our work. We did not use replicates in our exposure effect studies mainly due to costs. Second, we did not use factorial design for the SDCEO regimen. Thus, the SDCEO regimen will need to be studied in multiple extra clinical MAB isolates in the future, using factorial design. Third, our results on time-to-extinction were geared towards sterilization of MAB in cavities, which is more difficult test for the experimental regimens [2, 47, 49]. In tnodular disease, with lower *B_0_*, time-to-cure could be faster than in cavities [2, 47, 49].

To summarize, sulbactam-durlobactam-ceftriaxone trio (“double-double” peptidoglycan biogenesis inhibitors) combination led to faster time-to-extinction compared to sulbactam-durlobactam duo. A further addition of the two protein synthesis inhibitors in the SDCEO regimen suggested that the regimen could potentially shorten the therapy duration.

## Supporting information

Supplementary Methods and Results

## ACKNOWLEDGEMENTS

We thank Dr. Monika Kumaraswamy and Barbara Brown-Elliot, Mycobacteria & Nocardia Clinical Reference Laboratory, University of Texas Health Science at Tyler, Texas, for providing the clinical isolates. We also thank Prof. Rachel Thomson and Dr. Andrew Burke from the University of Queensland, Australia, for discussions on clinical translation of the findings.

## ETHICAL APPROVAL

Not applicable.

## DATA AVAILABILITY STATEMENT

Upon a reasonable request, the raw data for the results presented in the manuscript is available with the corresponding author.

## AUTHOR CONTRIBUTIONS

All named authors meet the International Committee of Medical Journal Editors (ICMJE) criteria for authorship for this article, take responsibility for the integrity of the work, and have given their approval for this version to be published. Conceptualization and design: Shashikant Srivastava (SKS) and Tawanda Gumbo (TG). Hollow Fiber model studies: Sanjay Singh (SS), GDB, and AS. Drug concentration measurement: MCL, BR. Data analysis, and PK/PD modeling: TG, SKS, SS. MCE for fixed-dose combinations: TG. Clinically relevant comments: PJM and TG. TG wrote the first draft of the manuscript. All authors read, edited, and approved the final version of the manuscript.

## ADDITIONAL INFORMATION

The Supplementary Figure and Supplementary Table are available online under “Supplementary Data”.

