## Supplementary Methods and Results for "Sulbactam-Durlobactam Plus Ceftriaxone Dosing and Novel Treatment Regimens for *Mycobacterium abscessus* Lung Disease"

### SUPPLEMENTARY DATA

Supplementary Methods, Results, and References

### SUPPLEMENTARY FIGURES

Supplementary Figure S1. Sulbactam concentrations observed in the HFS-MAB exposure-response studies.

Supplementary Figure S2. Durlobactam concentrations observed in the HFS-MAB exposure-response studies.

Supplementary Figure S3. Drug concentrations observed in the HFS-MAB inoculated with clinical isolates.

Supplementary Figure S4. Durlobactam probability of target attainment.

### SUPPLEMENTARY TABLES

Supplementary Table S1. Drug infusion in the HFS-MAB studies.

Supplementary Table S2. Study Quality Scoring.

Supplementary Table S3. Minimum inhibitory concentrations in 63 *Mycobacterium abscessus* isolates.

Supplementary Table S4. Model choice of PK/PD index based on corrected Akaike Information Criteria for microbial kill in the HFS-MAB.

Supplementary Table S5. Drug Minimum Inhibitory Concentrations mg/L Changes with Treatment.

Supplementary Table S6. Drug concentrations achieved in combination therapy studies.

Supplementary Table S7. Monte Carlo experiments domain of input versus output for normal renal function.

Supplementary Table S8. Dosing table for patients with MAB-LD with different creatinine clearances.

### SUPPLEMENTARY METHODS

#### Rationale for the testing double β-lactam combinations with a β-lactamase inhibitor, and choice of ceftriaxone as a companion drug.

First, as the rationale for the studies reported here, ceftriaxone and sulbactam both inactivate PonA1, PonA2, and D,D-c PbpA, durlobactam and sulbactam both inactivate $Bla_{Mab}$, while durlobactam inactivates all $Ldt_{Mab}$ (except $Ldt_{Mab3}$) and PonA1, which means extensive target redundancy [1]. In patients' lung cavities, the day 0 or pretreatment burden ($B_0$) ranges between 7.0-8.0 $\log_{10}$ CFU/lung, and is the major determinant of therapy outcomes, consistent with the *in vitro* inoculum effect [2-7]. Therefore, drugs for the target redundancy strategy must be chosen based on how extensively they kill MAB below $B_0$, which relegates cefuroxime, ceftaroline, and cefoxitin to the second tier [8, 9]. Another rationale for the choice of sulbactam-durlobactam is clinical pharmacokinetics (PK) and pharmacodynamics (PD); PK/PD is the scientific basis for optimized and practical dosing strategies [10]. Ceftriaxone has an 8h half-life, and sulbactam-durlobactam have half-life of 2.5h, which could lead to once or twice a day dosing, more practical than imipenem, which is dosed four to six times a day [11].

#### Minimal Inhibitory concentrations and time-kill studies

Stock cultures of the bacteria (*M. abscessus* ATCC#19977 and 63 clinical isolates) were grown to log phase cultures in Middlebrook 7H9 supplemented with 10% oleic acid-albumin-dextrose-catalase (OADC), followed by turbidity adjustment to McF 0.5 using a densitometer (DEN-1B, Grant-Bio) and back dilution to prepare the inoculum with an intended bacteria density of ~$10^5$

CFU/mL. We did not use cation-adjusted Mueller-Hinton broth (CAMHB), as CLSI recommended, as in our experience, MICs tend to be higher in that medium for the study drugs. We performed the following MIC experiments: (i) sulbactam-durlobactam MIC were performed over a concentration range of 0.25 mg/L to 128 mg/L alone and with 256 mg/L fixed concentration of ceftriaxone, (ii) ceftriaxone MICs over a concentration range of 0.25 mg/L to 128 mg/L alone and with 4 mg/L fixed concentration of sulbactam-durlobactam, (iii) MIC of all other study drugs alone without combination of sulbactam-durlobactam or ceftriaxone. All the MIC experiments were performed twice in duplicate. MICs were recorded after 72 hours of incubation at $30^0$C. Three different individuals read the MICs, and in case of trailing effect, agreement between two operators was recorded as the MIC value.

For the static time-kill studies, inoculum preparation for the ATCC strain was the same as described above. The concentration of each drug used in the experiments, alone and as various combinations with sulbactam-durlobactam, is listed in **Supplementary Table S1**, except azithromycin (0.65 mg/L peak concentration), which was used only in the time-kill studies. Experiments were carried out in a total volume of 5 mL, in triplicate for each condition. Cultures were co-incubated with drug(s) for 72 hours of incubation at $30^0$C. Next, 1 mL cultures were collected in sterile 1.8 mL centrifuge tubes, washed twice with normal saline by centrifugation at 13,000 rpm for 5 min at ambient temperature. Bacterial pellets were resuspended in 1 mL normal saline, 10-fold serially diluted, and spread on Middlebrook 7H10 agar for CFU estimation.

**HFS-MAB exposure-effect**

The construction and design of the HFS-MAB have been extensively described in the literature [8, 9, 11-16]. Here, we combined an exposure-effect study with a dose-scheduling design, using the ATCC#19977 strain. Drugs were infused to mimic the intrapulmonary PKs following human equivalent doses of the commercial 1-to-1 sulbactam and durlobactam for injection, 0, 3G every 24h (q24h), 1.5G q12h, 1G q8h, 4.5G q24h, 3G q12h, 1.5G q8h, 6G q24h, 3G q12h, 2G q8h, and

3G q8h. The drug exposures shown in the **Supplementary Table S1** were delivered via programmable syringe pumps. The central compartment of each HFS-MAB unit was sampled for drug concentration measurements at pre-dose and 3, 5, 8, 11, 16, 19, and 23.5h post-dosing. The peripheral compartments were sampled for bacterial burden on days 0, 1, 2, 3, 4, 7, 10, and 14.

**Drug concentration measurement**

We used the previously published methods for ceftriaxone, epetraborole, omadacycline, and minocycline concentration measurements in the HFS-MAB samples [16-19]. The following is a brief description of the methods we developed for the measurement of sulbactam and durlobactam in the HFS-MAB samples. Sulbactam, durlobactam, and internal standards were purchased from Toronto Research Chemicals (LGC), Toronto. All other chemicals were chromatographic or LC-MS/MS grade and purchased from Fisher Scientific, USA. We employed stable-isotope dilution liquid chromatography-electrospray ionization tandem mass spectrometry (LC-MS/MS) to determine the analytes in the experimental samples (matrix Middlebrook 7H9 broth supplemented with 2% dextrose). LC-MS/MS analysis was performed using Waters Acquity UPLC connected to a Waters Xevo TQD mass spectrometer (Milford, MA). Data was collected using MassLynx version 4.2 SCN985 software. Separation was achieved on a Waters Acquity UPLC BEH C18 1.7 µm 50 x 2.1mm analytical column. All standard and internal standard (IS) stock solutions were prepared at 1 mg/ml in methanol and stored at -20 C. A 7-point calibration curve, low-, mid-, and mid high and high-quality control samples (QCL QCM, QC MH and QCH) were prepared by diluting the stock solution in 50% MeOH and water. In a 96-well plate, 20µl of SC or QC was added to 180µL of 7H9 + 2% dextrose broth or 200 µl of sample were added to 20 µl of IS solution and 400 µl of CLR water and mixed. The separation gradient used mobile phase A (a mixture of aqueous FA/ammonium formate) and mobile phase B was methanol. The flow rate was 0.6 mL/min with a total run time was 3.5 mins. Compounds were detected using ESI in MRM mode. The mass charge ratio for sulbactam, sulbactam-d2, durlobactam, and azithromycin-d3, sulbactam were 232 > 140, 234 > 190, 276 > 96.8, and 752.2 > 158, respectively. The standard curve ranged from 100 to 20000 ng/mL. The LLOQ was 100 ng/mL, and the low, mid, mid-high-, and high-quality control (QC) concentrations were 300, 3000, 8000, and 16,000 ng/mL. Validation performance showed %CV and %E below 20%. Run QC criteria specified that 5/8 QC (L, M, MH, and H) had to have less than 20% or less error and no two over 20% at the same concentration.

**Monte Carlo experiments (MCE)**

MCE were performed as described in detail in our past publication. In patients being treated for Gram-negative bacilli the drug 1G of sulbactam and 1G of durlobactam are infused over 3h every 4-6h, for a duration of treatment of 7 to 14 days [20]. Both sulbactam and durlobactam are two-compartment model drugs, based on Cammarata et al [21]. The epithelial lining fluid (ELF)-to-plasma penetration relative for durlobactam was 37.2% while that for sulbactam was 53.3%, in population pharmacokinetic analyses [21]. Since 75% to 85% of sulbactam and durlobactam are excreted unchanged in urine, the most important covariate for clearance is renal function. Therefore, we examined PTA and cumulative fraction of response of each dose by creatinine clearance in 4 categories: (1) >90 mL/min which is normal renal function, (2) 60-90 mL/min which is mild renal dysfunction,(3) ≥30 to <60 mL/min moderate renal dysfunction, and (4) <30 mL/min which is severe. Since the dosing of the drug every 4h for many months would impose a hardship, we only examined q8h and q12h dosing schedules. We examined for probability of target attainment (PTA) for 1G each of sulbactam and durlobactam, or 2G, or 3G, or 4G either twice each day or 3 times each day, infused over 3h, at each MIC. We examined the ability of each of these doses to achieve or exceed $EC_{80}$ in the lungs of 10,000 virtual subjects (90,000 subjects total). Since protein binding is 10% for durlobactam and 38% for sulbactam, and the mean protein ELF/plasma ratios are 0.13-0.25 for total protein and 0.1-0.19 for albumin, the effect of protein binding in ELF is expected to be negligible [22-25]. Thus, we used the total concentrations measured in ELF by Cammarata et al [21].

### SUPPLEMENTARY RESULTS

#### Exposure-response HFS-MAB PKs

The sulbactam concentrations measured in the HFS-MAB units are shown in **Supplementary Figure S1A-C**. PK modeling revealed a sulbactam clearance of 6.93 $x10^{-3}$ L/h, a volume of 47.19 $x10^{-3}$ L. The observed versus model-predicted concentrations are shown in **Figure S1D**. The durlobactam concentrations measured in the HFS-MAB units are shown in **Supplementary Figure S2A-C**. PK modeling revealed a durlobactam clearance of 6.56 $x10^{-3}$ L/h, a volume of 47.71 $x10^{-3}$ L. The observed versus model-predicted concentrations are shown in **Figure S2D**. The PK modeled data for sulbactam and durlobactam were used to calculate AUC/MIC ratios, peak concentration to MIC ratio, and time concentration persists above MIC ($\%T_{MIC}$); the MAB ATCC#19977 inoculated in HFS-MAB had an MIC of 4 mg/L for sulbactam-durlobactam (1:1 combination).

#### Durlobactam probability of target attainment (PTA)

The PTAs for durlobactam for the target exposure of $\%T_{MIC}$ of 10% in the lungs, were as shown in **Supplementary Figure S4**. In **Figure S4A**, normal renal function, the lowest dose of 1G every 24h achieved PTA below 90% at an MIC of 4 mg/L. In **Figure S4B** with a creatinine clearance of 60-90 mL/min, 1G every 24h achieved PTA below 90% at an MIC of 8 mg/L. In **Figure S4C**, for a creatinine clearance of ≥30 to <60 mL/min, the same dose fell below PTA of 90% at an MIC of 8 mg/L. In severe renal dysfunction in **Figure S4D**, with creatinine clearance less than 30 ml/min, 1G every 24h achieved PTA below 90% at an MIC of 16 mg/L.

#### Resistance in combination therapy studies

If resistance is taken as >1-tube dilution increase in MIC compared to day 0, then sulbactam-durlobactam duo (R1) and sulbactam-durlobactam-ceftriaxone (R3) consistently failed to prevent acquired AMR. This means that sulbactam-durlobactam duo and the sulbactam-durlobactam-ceftriaxone trio need other drugs to suppress AMR.

**Supplementary Table S1. Drug infusion in the HFS-MAB studies.**

| | Frequency (h) | Infusion time (h) | Half-life (h) | $AUC_{0-24}$ (mg*h/L) | Peak concentration (mg/L) | $T_{MIC}$ % |
|---|---|---|---|---|---|---|
| **Exposure-effect and dose fractionation** | | | | | | |
| R1 | 0 | 0 | 0 | 0 | 0 | 0 |
| R2 | 0 | 0 | 0 | 0 | 0 | 0 |
| R3 | 24 | 3 | 2.5 | 45 | 16 | 13 |
| R4 | 24 | 3 | 2.5 | 279 | 96 | 33 |
| R5 | 12 | 3 | 2.5 | 274 | 48 | 50 |
| R6 | 8 | 3 | 2.5 | 195 | 32 | 63 |
| R7 | 12 | 3 | 2.5 | 411 | 72 | 58 |
| R8 | 8 | 3 | 2.5 | 390 | 48 | 75 |
| R9 | 24 | 3 | 2.5 | 558 | 192 | 42 |
| R10 | 12 | 3 | 2.5 | 539 | 96 | 67 |
| R11 | 8 | 3 | 2.5 | 767 | 64 | 79 |
| R12 | 8 | 3 | 2.5 | 780 | 96 | 100 |
| **Combination therapy** | | | | | | |
| Sulbactam/durlobactam | 12 | 3 | 2.5 | 407 | 72 | >50% |
| Ceftriaxone | 24 | 1 | 8 | 2710 | 260 | 100 |
| Minocycline | 24 | 1 | 15 | 50 | 3.5 | |
| Omadacycline | 24 | 1 | 15 | 24 | 1.8 | |
| Epetraborole | 12 | 1 | 11 | 47.5 | 3.4 | |

**Supplementary Table S2. Minimum inhibitory concentrations in 63 *Mycobacterium abscessus* isolates.**

| Isolate ID | SUL/DUR | SUL/DUR + CTX 256 mg/L | Tube dilution change | CTX | CTX+SUL/DUR (4 mg/L) | Tube dilution change |
|---|---|---|---|---|---|---|
| **ATCC#19977** | 4 | 0.25 | 4 | 256 | 0.5 | 9 |
| Mab_1 | 8 | <0.125 | 7 | 256 | 0.5(T) | 9 |
| Mab_2 | 8 | <0.125 | 7 | 256 | 2 | 7 |
| Mab_3 | 8 | <0.125 | 7 | 256 | 0.5 | 9 |
| Mab_4 | 8 | <0.125 | 7 | 256 | 0.5 | 9 |
| Mab_5 | 8 | <0.125 | 7 | 256 | 2 | 7 |
| Mab_6 | 8 | <0.125 | 7 | 256 | 1 | 8 |
| Mab_7 | 8 | <0.125 | 7 | 256 | 1 | 8 |
| Mab_8 | 8 | <0.125 | 7 | 256 | 2 | 7 |
| Mab_9 | 8 | <0.125 | 7 | 256 | 16 | 4 |
| Mab_10 | >64 | 64 | | 256 | 64 | 2 |
| Mab_11 | 16 | <0.125 | 8 | 256 | 2 | 7 |
| Mab_12 | 16 | <0.125 | 8 | 256 | 8 | 3 |
| Mab_13 | 32 | <0.125 | 9 | 256 | 1 | 8 |
| Mab_14 | 32 | <0.125 | 9 | 256 | ≤0.5 | 10 |
| Mab_15 | 32 | <0.125 | 9 | 256 | 1 | 8 |
| Mab_16 | 32 | <0.125 | 9 | 256 | 1 | 8 |
| Mab_17 | 32 | <0.125 | 9 | 256 | 1 | 8 |
| Mab_18 | 32 | <0.125 | 9 | 256 | 1 | 8 |
| Mab_19 | 8 | <0.125 | 7 | 256 | ≤0.5 | 10 |
| Mab_20 | 8 | <0.125 | 7 | 256 | 32 | 3 |
| Mab_21 | 8 | 0.25 | 7 | 256 | 0.5 | 9 |
| Mab_22 | 64 | 16 | 2 | 256 | 64 | 2 |
| Mab_23 | 64 | 8 | 3 | 256 | 16 | 4 |
| Mab_24 | 32 | <0.125 | 9 | 256 | 32 | 3 |
| Mab_25 | 32 | 1 | 5 | 256 | 32 | 3 |
| Mab_26 | 4 | 0.25 | 4 | 256 | 32 | 3 |
| Mab_27 | 64 | 0.25 | 8 | 256 | 16 | 4 |
| Mab_28 | 64 | 16 | 2 | 256 | 256 | 0 |
| Mab_29 | 64 | 1 | 6 | >256 | 256 | 0 |
| Mab_30 | 64 | 0.25 | 8 | 256 | 128 | 1 |
| Mab_31 | 32 | 0.5 | 6 | 256 | 8 | 3 |
| Mab_32 | 16 | 0.25 | 6 | 256 | 64 | 2 |
| Mab_33 | 32 | <0.125 | 9 | 256 | 128 | 1 |
| Mab_34 | 32 | 0.5 | 8 | 256 | 16 | 4 |
| Mab_35 | 64 | <0.125 | 10 | 256 | 8 | 5 |

| | | | | | | |
|---|---|---|---|---|---|---|
| Mab_36 | 32 | 1 | 5 | 256 | 8 | 5 |
| Mab_37 | 32 | <0.125 | 9 | 256 | 8 | 5 |
| Mab_38 | 32 | <0.125 | 9 | 256 | 4 | 6 |
| Mab_39 | 32 | <0.125 | 9 | 256 | 8 | 5 |
| Mab_40 | 64 | <0.125 | 9 | 256 | 32 | 3 |
| Mab_41 | 64 | <0.125 | 9 | 256 | 16 | 4 |
| Mab_42 | 8 | <0.125 | 9 | 256 | 128 | 1 |
| Mab_43 | 64 | 8 | 3 | 256 | 64 | 2 |
| Mab_44 | 128 | 0.25 | 9 | 256 | 64 | 2 |
| Mab_45 | 128 | 0.5 | 8 | 256 | 64 | 2 |
| Mab_46 | 64 | <0.125 | 9 | 256 | 16 | 4 |
| Mab_47 | 128 | 8 | 4 | 256 | 64 | 2 |
| Mab_48 | 64 | <0.125 | 9 | 256 | 128 | 1 |
| Mab_49 | 64 | <0.125 | 9 | 256 | 128 | 1 |
| Mab_50 | 128 | <0.125 | 10 | 256 | 128 | 1 |
| Mab_51 | 16 | <0.125 | 8 | 256 | 32 | 3 |
| Mab_52 | 32 | <0.125 | 9 | 256 | 16 | 4 |
| Mab_53 | 32 | <0.125 | 9 | 256 | 64 | 2 |
| Mab_54 | 32 | <0.125 | 9 | 256 | 64 | 2 |
| Mab_55 | 32 | 0.5 | 8 | 256 | 64 | 2 |
| Mab_56 | 32 | <0.125 | 9 | 256 | 256 | 0 |
| Mab_57 | 32 | <0.125 | 9 | 256 | 256 | 0 |
| Mab_58 | 16 | <0.125 | 8 | 128 | 128 | 0 |
| Mab_59 | 32 | <0.125 | 9 | 256 | 64 | 2 |
| #8586/BO | NA | NA | NA | 256 | <0.5 | 10 |
| #9152 | 8 | 2 | 2 | 256 | 128 | 1 |
| #7630 (Smooth) | 4 | 0.25 | 4 | 256 | 256 | 0 |
| #9694 | >64 | 32 | 2 | 256 | 256 | 0 |

SUL, sulbactam; DUR, durlobactam; CTX, ceftriaxone; NA, not available.

**Supplementary Table S3. Model choice of PK/PD index based on corrected Akaike Information Criteria for microbial kill in the HFS-MAB.**

| | **PK/PD parameter** | **Day 1** | **Day 2** | **Day 3** | **Day 4** | **Day 7** | **Day 10** | **Day 14** |
|---|---|---|---|---|---|---|---|---|
| **Sulbactam** | $\%T_{MIC}$ | **-20.20** | **-13.05** | **9.77** | **5.84** | **1.35** | **-1.68** | **-2.15** |
| | AUC/MIC | -11.52 | -13.04 | 9.86 | 5.95 | 2.05 | -1.29 | 3.92 |
| | $C_{max}$/MIC | 8.63 | -13.04 | 10.33 | 6.47 | 2.62 | 0.46 | 5.82 |
| **Durlobactam** | | | | | | | | |
| | $\%T_{MIC}$ | **-9.92** | -13.04 | **10.33** | **5.78** | **2.62** | **0.46** | **5.93** |
| | AUC/MIC | **-9.92** | -13.04 | **10.33** | 6.28 | **2.62** | 1.57 | **5.93** |
| | $C_{max}$/MIC | **-9.92** | **-13.07** | **10.33** | 5.79 | **2.62** | 0.47 | 11.57 |

PK/PD, pharmacokinetics/pharmacodynamics

**Supplementary Table S4. Drug minimum inhibitory concentrations (MIC in mg/L) change with treatment in HFS-MAB.**

| | **Day 0** | **Day 14** | | | |
|---|---|---|---|---|---|
| **REGIMEN (R)** | **Inoculum** | **Sulbactam-durlobactam (R1)** | **Ceftriaxone (R2)** | **Sulbactam-durlobactam-Ceftriaxone (R3)** | **Non-treated (R4)** |
| **Isolate #1 (MAB_2)** | | | | | |
| Sulbactam-durlobactam | 8 | 16 | 8 | 16 | 8 |
| Ceftriaxone | >256 | >256 | >256 | >256 | >256 |
| Sulbactam-durlobactam +256 mg/L ceftriaxone | ≤0.125 | ≤0.25 | ≤0.25 | ≤0.25 | ≤0.25 |
| Ceftriaxone + 4 mg/L sulbactam-durlobactam | 0.5 | **16** | **8** | **32** | **8** |
| **Isolate #2 (MAB_12)*** | | | | | |
| Sulbactam-durlobactam | 16 | 16 | 16 | **128** | 16 |
| Ceftriaxone | >256 | >256 | >256 | >256 | >256 |
| Sulbactam-durlobactam +256 mg/L ceftriaxone | <0.125 | **0.5** | ≤0.25 | **32** | ≤0.25 |
| Ceftriaxone + 4 mg/L sulbactam-durlobactam | 8 | **64** | 16 | **256** | 16 |

*The same isolate was used in the two-drug and multi-drug combination HFS-MAB studies.

**Supplementary Table S5. Drug Concentrations achieved in combination therapy HFS-MAB studies.**

| | $AUC_{0-24}$ (mg*h/L) | $C_{max}$ (mg/L) | $\%T_{MIC}$ |
|---|---|---|---|
| **Sulbactam** | 652.5 ± 61.18 | 46.29 | 62.5% |
| **Durlobactam** | 867.1 ± 142.1 | 65.15 | 62.5% |
| **Ceftriaxone** | 5,558 ± 1,305 | 314.1 | 50% |
| **Epetraborole** | 53.89 ± 1.70 | 3.33 | 100% |
| **Minocycline** | 44.6 ± 1.47 | 3.24 | 0% |
| **Omadacycline** | 27.34 ± 1.19 | 1.82 | 95.83% |

**Supplementary Table S6. Monte Carlo experiments domain of input versus output for normal renal function.**

| | Subroutine PRIOR | | Virtual subjects | |
|---|---|---|---|---|
| | **Estimate** | **%CV** | **Estimate** | **%CV** |
| **Sulbactam** | | | | |
| Clearance (L/h) | 13.5 | 47.0 | 13.50 | 46.51 |
| Central volume (L) | 12 | 31.1 | 12.09 | 31.08 |
| Distributional clearance (L/h) | 7.88 | 25.8 | 7.97 | 25.81 |
| Peripheral Volume (L) | 6.99 | 44.3 | 6.92 | 44.05 |
| **Durlobactam** | | | | |
| Clearance (L/h) | 9.33 | 27.9 | 9.32 | 27.9 |
| Central volume (L) | 12.5 | 27.5 | 12.46 | 28.22 |
| Distributional clearance (L/h) | 4.43 | 40 | 4.42 | 40.22 |
| Peripheral Volume (L) | 5.83 | 27.8 | 5.84 | 27.8 |

**Table S7. Dosing table for patients with MAB-LD with different creatinine clearances.**

| Creatinine clearance in mL/min | Dose (grams) | Frequency (hours) |
|---|---|---|
| >90 | 2 | 8 |
| 60-90 | 1 | 8 |
| ≥30 to <60 | 1 | 12 |
| <30 | 1 | 24 |

**Supplementary Figure S1. Sulbactam concentrations observed in the HFS-MAB exposure-response studies.**

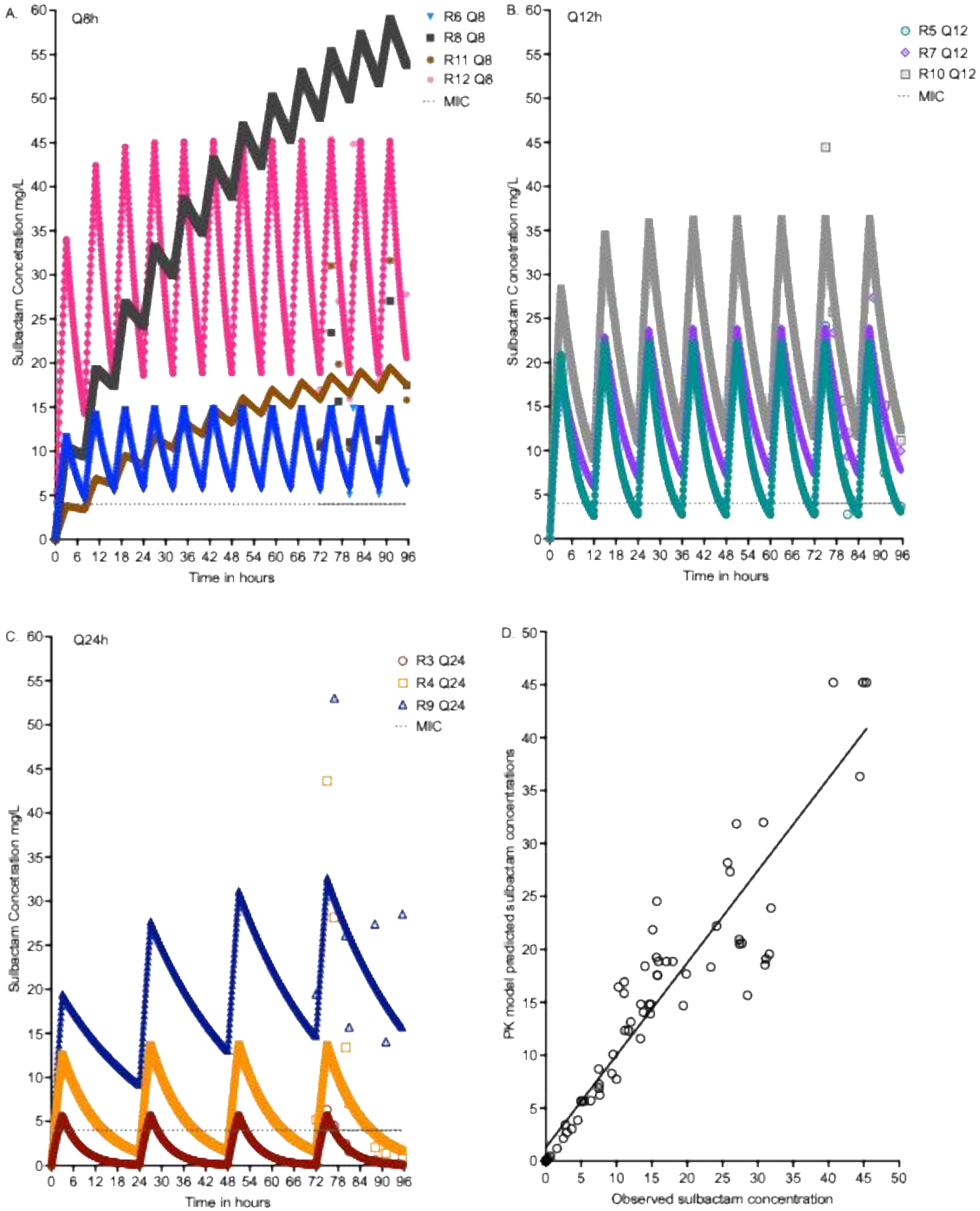

**Supplementary Figure S2. Durlobactam concentrations observed in the HFS-MAB exposure-response studies.**

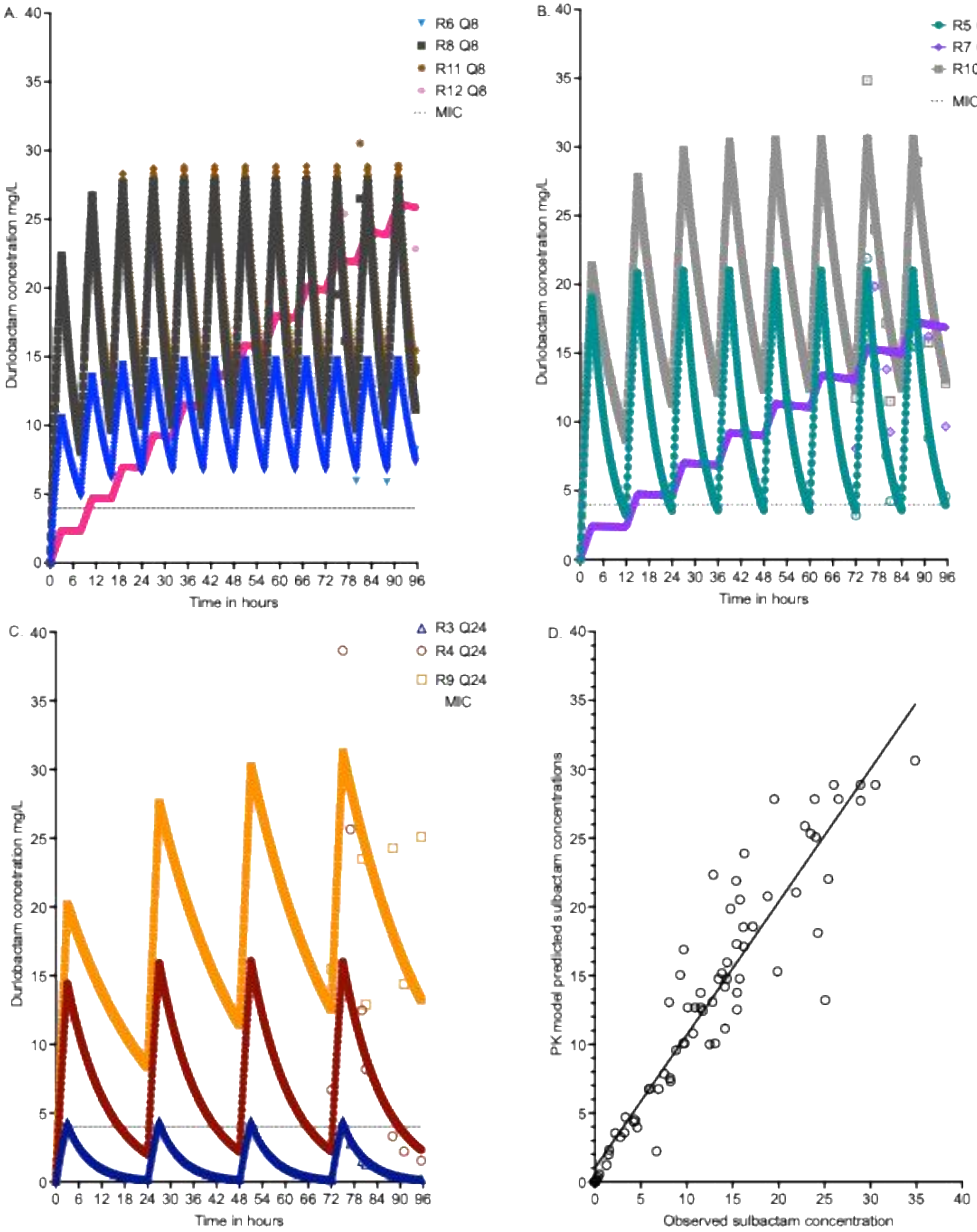

**Supplementary Figure S3. Drug concentrations observed in the HFS-MAB inoculated with the clinical isolates.**

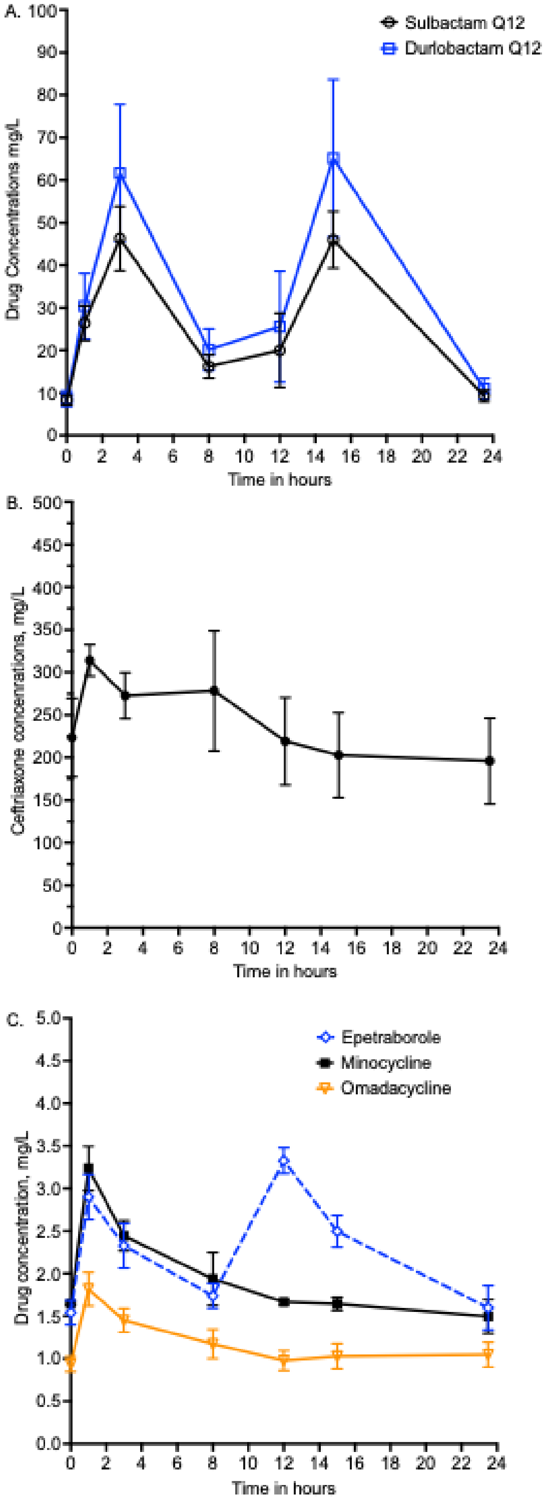

**Figure S4. Durlobactam probability of target attainment.**

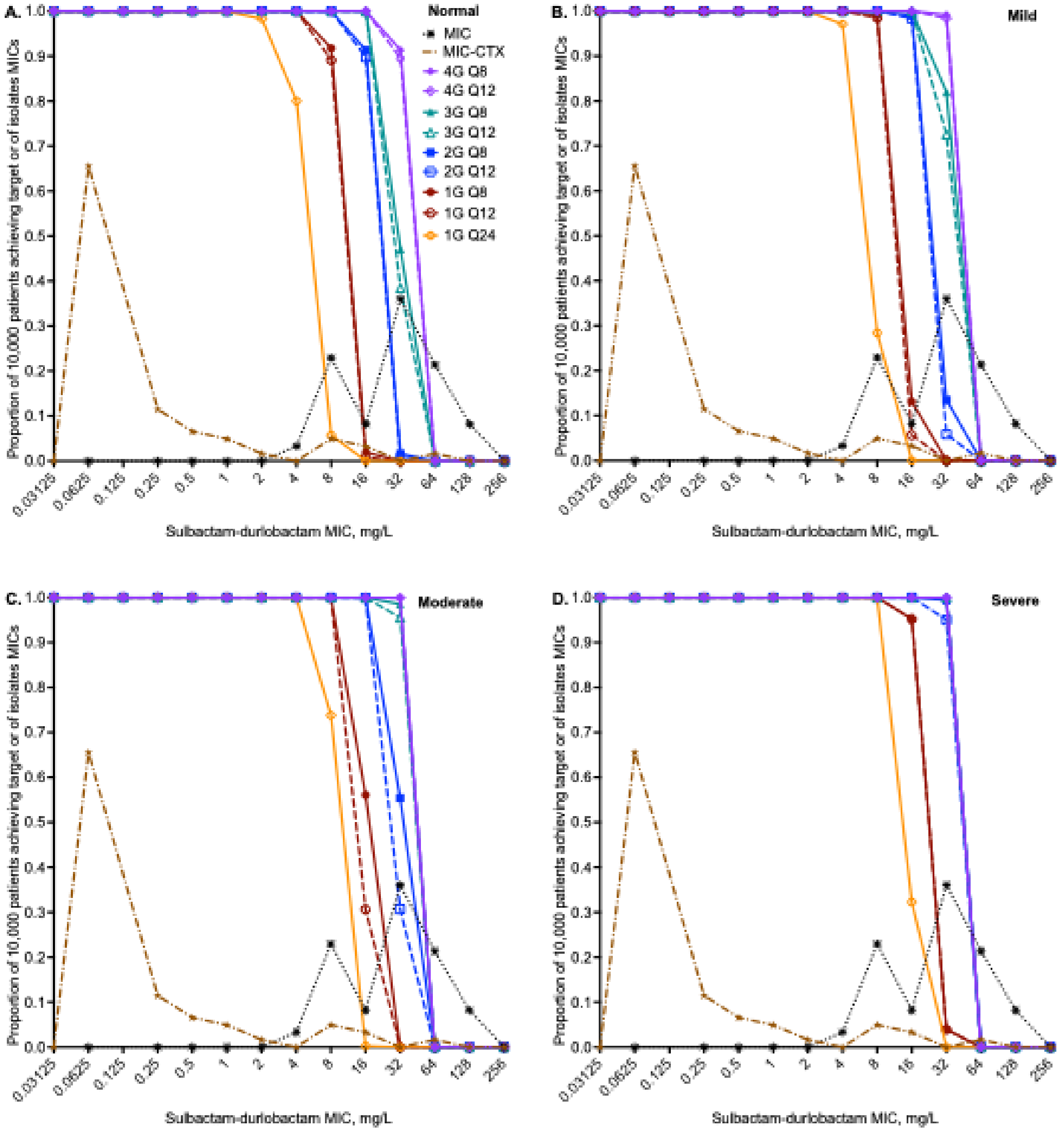
